# Fluorescence Microscopy Datasets for Training Deep Neural Networks

**DOI:** 10.1101/2020.06.17.158097

**Authors:** Guy M. Hagen, Justin Bendesky, Rosa Machado, Tram-Anh Nguyen, Tanmay Kumar, Jonathan Ventura

## Abstract

**Background:** Fluorescence microscopy is an important technique in many areas of biological research. Two factors which limit the usefulness and performance of fluorescence microscopy are photobleaching of fluorescent probes during imaging, and, when imaging live cells, phototoxicity caused by light exposure. Recently developed methods in machine learning are able to greatly improve the signal to noise ratio of acquired images. This allows researchers to record images with much shorter exposure times, which in turn minimizes photobleaching and phototoxicity by reducing the dose of light reaching the sample.

**Findings:** To employ deep learning methods, a large amount of data is needed to train the underlying convolutional neural network. One way to do this involves use of pairs of fluorescence microscopy images acquired with long and short exposure times. We provide high quality data sets which can be used to train and evaluate deep learning methods under development.

**Conclusion:** The availability of high quality data is vital for training convolutional neural networks which are used in current machine learning approaches.

## Data description

### Context

Fluorescence microscopy is an important technique in many areas of biomedical research, but its use can be limited by photobleaching of fluorescent probe molecules caused by the excitation light which is used. In addition, reactive oxygen species which are generated by exposing samples to light can cause cell damage and even cell death, limiting imaging of live cells [1,2]. Many strategies have been devised to overcome this problem including the use of specialized culture media [3,4], pulsed excitation [5], or more elaborate methods such as controlled light exposure microscopy [6,7].

Another approach involves recording of fluorescence microscopy images with short exposure times, low excitation light intensity, or both. This results in images with low signal to noise ratios (SNRs), which can subsequently be improved using a variety of image restoration approaches [8–12]. Noise in low light images of this type typically follows a Poisson-Gaussian distribution. This condition makes solving the inverse problem which arises in image restoration methods very difficult, leading to a variety of approximate methods [13].

Recently, deep learning methods in artificial intelligence [14] have been applied to many problems in image analysis, including those in optical microscopy [15–18] and in image denoising [19,20]. Deep learning approaches typically require a large amount of data to train the underlying convolutional neural network [21], however, such datasets are not always available. Here we provide fluorescence microscopy datasets which can be used to train and evaluate neural networks for the purpose of image denoising. The dataset consists of pairs of images acquired with different exposure times. After training, the network can subsequently be used to enhance the SNR of newly acquired images.

One advantage of deep learning methods is that they can learn a task such as denoising from the data itself, thus providing a sample-specific method which does not depend on a physical model. Once a network has been trained, subsequent image denoising using a convolutional neural network is fast compared to traditional methods which are typically much slower.

In addition to providing the datasets, we evaluated the performance of a recently proposed neural network for content-aware image restoration (CARE) of fluorescence microscopy images [17]. To do this we used CSBDeep [22], a toolbox for implementation of the CARE network. This network uses a series of convolutional layers in a U-Net architecture [23].

We also evaluated a self-supervised learning approach called a blindspot neural network [24], an extension of the Noise2Void approach [25]. This method learns denoising using only the noisy data. It uses a U-Net style architecture but uses careful padding and cropping to force the network to learn to predict the value of each denoised pixel based on the neighborhood of that pixel in the noisy input. We used our own implementation in Python using the Keras library.

## Methods

### Fluorescence Microscopy

We acquired datasets 1 - 3 using an IX83 microscope equipped with UplanSApo 60×/1.3 NA oil immersion and UplanSApo 20×/0.75 NA air objectives (Olympus, Tokyo, Japan), Zyla 4.2-plus sCMOS camera (Andor, Belfast, UK), and SpectraX light source (Lumencor, Beaverton, OR, USA). Focusing was achieved using a piezo-Z stage (Applied Scientific Instrumentation, Eugene, OR, USA). The system was controlled by IQ3 software (Andor). We used fluorescence filter set 59022 (Chroma, Bellows Falls, VT, USA). Dataset 4 was acquired with a SP5 laser scanning confocal microscope (Leica, Mannheim, Germany) using 488 nm and 543 nm lasers and a HCX PL APO CS 63×/1.4 NA oil immersion objective (Leica).

The sample in all cases was a FluoCells #1 prepared slide (Molecular Probes, Eugene, OR, USA). This slide contains bovine pulmonary artery endothelial cells stained with MitoTracker Red CMXRos (labels mitochondria) and AlexaFluor 488 phalloidin (labels actin).

### Data Analysis

Each dataset consisted of images of size 2048×2048 pixels, where the last ten percent were used for testing and the remaining were used for training. In datasets 1, 2, and 3 we had 100 image pairs in each dataset and in dataset 4 (confocal), we had 79 image pairs and the image size was 1024×1024 pixels.

To train the CARE network, we used the following configuration. We used the ADAM optimizer [26], the training batch size was 16 images, the number of training epochs was 200, the initial learning rate was 0.0004, and the iterations per epoch (training steps) was 400. In sampling the training images, 800 patches per image of size 64 pixels by 64 pixels were used to train the CARE network. In all experiments, images were split according to the ratio 5:1 for training and validation respectively.

Following the standard implementation of the CSBDeep network, we used the Laplacian loss function

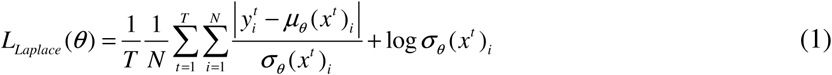

where T is the number of training images, N is the number of pixels per image, *y*^*t*^ is the ground truth pixel value, *x*^*t*^ is the input pixel, *μ* and *σ* are the mean and variance of the predicted pixel distribution.

To train the blindspot network, we used the same configuration described in Laine, et al (2019). We used the ADAM optimizer and trained on random crops of size 128×128 pixels using a mean-squared-error loss function. We trained over 200 epochs with 400 steps per epoch and an initial learning rate of 0.0003. In all experiments, the last five images of the training set were withheld for validation. For comparison we used a standard denoising method, block matching and 3D filtering (BM3D) [27].

## Results

We acquired four datasets under different conditions. Table 1 provides an overview of the four datasets. In the widefield data, we used the Lumencor light source control software to adjust the illumination intensity such that the desired signal to noise levels were achieved.

**Table 1:** Overview of the datasets.

| data set | 1 (60X noise level 1) | 2 (60X noise level 2) | 3 (20X) | 4 (confocal) |
| --- | --- | --- | --- | --- |
| microscope | Widefield | Widefield | Widefield | Confocal |
| objective | 60×/1.35NA oil immersion | 60×/1.35NA oil immersion | 20×/0.75NA air | 63×/1.4NA oil immersion |
| pixel size | 108 nm | 108 nm | 325 nm | 96 nm |
| Exposure times | high exposure (actin): 400 ms | high exposure (actin): 1000 ms | high exposure (actin): 500 ms | 0.7 images/sec |
|  | low exposure (actin): 20 ms | low exposure (actin): 15 ms | low exposure (actin): 20 ms | 0.7 images/sec |
|  | high exposure (mitochondria): 400 ms | high exposure (mitochondria): 600 ms | high exposure (mitochondria): 400 ms | 0.7 images/sec |
|  | low exposure (mitochondria): 20 ms | low exposure (mitochondria): 10 ms | low exposure (mitochondria): 15 ms | 0.7 images/sec |
| image size, pixels | 2048 × 2048 | 2048 × 2048 | 2048 × 2048 | 1024 × 1024 |

After data acquisition, we tested three different methods for image denoising. Figure 1 shows the original low exposure image (raw), the matching high exposure image (ground truth), and the results of the CARE method, the blindspot method, and a standard denoising method (BM3D). For this comparison we selected an image pair from data set 1 (60X noise level 1).

**Figure 1:**
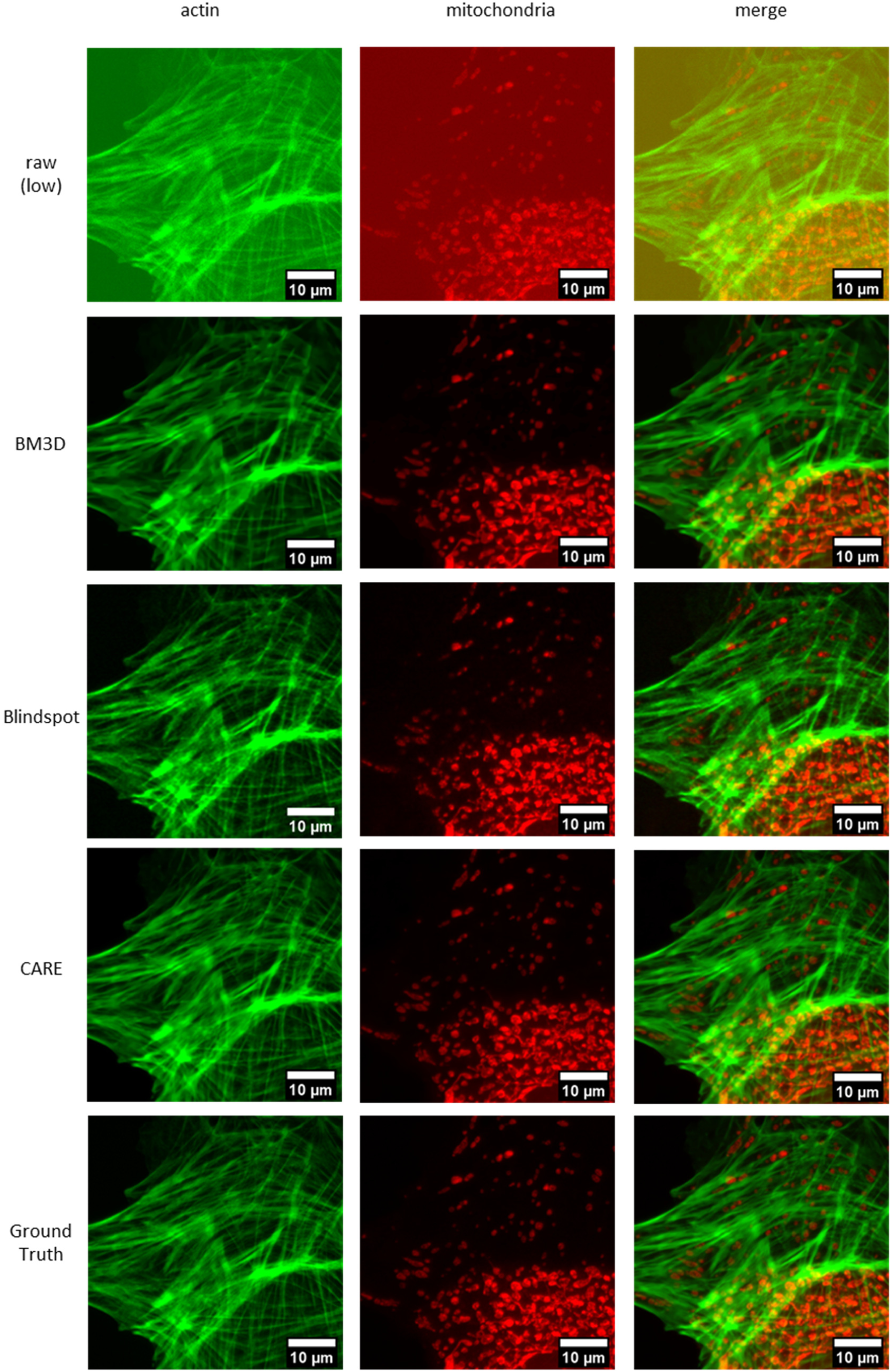
Results of denoising methods. Shown are selected images from dataset 1 (60X noise 1).

Table 2 provides average metrics for the denoising performance for each method on each dataset. We used two metrics: peak signal-to-noise ratio (PSNR) and structural similarity (SSIM). We normalized both images by clipping values below the 1^st^ percentile and above the 99^th^ percentile. We then scaled and shifted both images to minimize the mean squared error (MSE) between them [17].

**Table 2.**
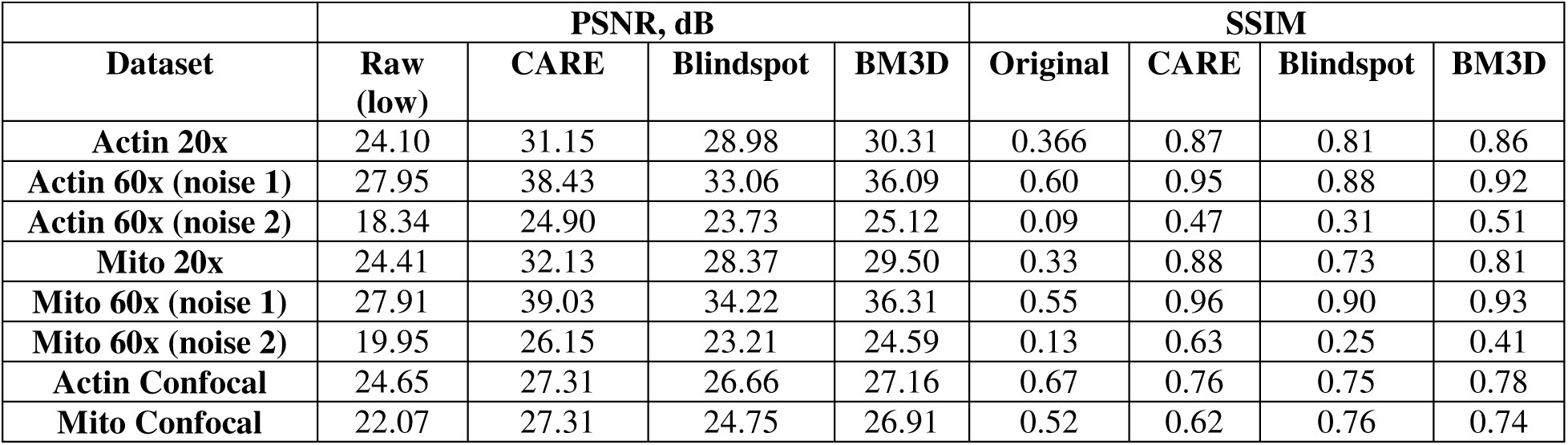
Average PSNR and SSIM results (n=10)

| Dataset | PSNR, dB |  |  |  | SSIM |  |  |  |
| --- | --- | --- | --- | --- | --- | --- | --- | --- |
|  | Raw (low) | CARE | Blindspot | BM3D | Original | CARE | Blindspot | BM3D |
| Actin 20x | 24.10 | 31.15 | 28.98 | 30.31 | 0.366 | 0.87 | 0.81 | 0.86 |
| Actin 60x (noise 1) | 27.95 | 38.43 | 33.06 | 36.09 | 0.60 | 0.95 | 0.88 | 0.92 |
| Actin 60x (noise 2) | 18.34 | 24.90 | 23.73 | 25.12 | 0.09 | 0.47 | 0.31 | 0.51 |
| Mito 20x | 24.41 | 32.13 | 28.37 | 29.50 | 0.33 | 0.88 | 0.73 | 0.81 |
| Mito 60x (noise 1) | 27.91 | 39.03 | 34.22 | 36.31 | 0.55 | 0.96 | 0.90 | 0.93 |
| Mito 60x (noise 2) | 19.95 | 26.15 | 23.21 | 24.59 | 0.13 | 0.63 | 0.25 | 0.41 |
| Actin Confocal | 24.65 | 27.31 | 26.66 | 27.16 | 0.67 | 0.76 | 0.75 | 0.78 |
| Mito Confocal | 22.07 | 27.31 | 24.75 | 26.91 | 0.52 | 0.62 | 0.76 | 0.74 |

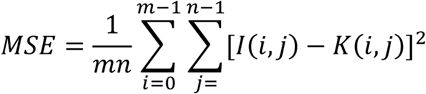

Where *I* is a high SNR image and *K* is the corresponding low SNR image after restoration. Finally, the PSNR metric was calculated as

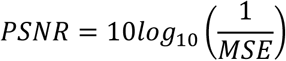

The SSIM metric [28] is an image quality metric designed to approximate human perception of similarity to a reference image. Unlike PSNR, the metric takes into account structural information in the image. The SSIM metric ranges from 0 to 1 with a greater number indicating higher quality.

As shown in Table 2, the unsupervised blindspot method is the weakest performer on both metrics. BM3D is better on both metrics but surpassed by the supervised CARE method on almost all datasets. All methods exhibit an approximately 10 dB drop in PSNR or greater on the noisier datasets (Noise 2) in comparison to Noise 1. Each method also performed about 6-7 dB worse on 20× magnification data in comparison to the 60× magnification data.

Visual inspection of the restored images (example shown in Figure 1) shows that, despite having high SSIM scores, the BM3D tends to blur the images more than the other methods. The results of the blindspot method are noticeably noisier than the results of the other methods.

Table 3 presents an comparison of the methods in terms of computation time. Using a single Nvidia V100 GPU, the CARE network took about 35 minutes to train on a single dataset while the blindspot network took about 2.6 hours. The CARE network took about 1 second to process a single image while the blindspot network took over 3 seconds. The BM3D method does not require training but took about 50 seconds to process a single image in MATLAB on a 2.6 GHz Intel Core i3-7100U processor.

**Table 3.** Training and processing times.

|  | Training time, sec | Processing time for 1 image, sec |
| --- | --- | --- |
| CARE | 2143.01 | 0.90 |
| Blindspot | 9450.86 | 3.35 |
| BM3D | - | 50 |

## Reuse potential

The provided data can be used to implement new methods in machine learning or to test modifications of existing approaches. The data can be used to evaluate methods for denoising, super-resolution, or generative modeling, as well as new image quality metrics, for example. The data could also be used to evaluate the generalization ability of methods trained on one type of data and tested on another. High quality, publicly available data of this type has been lacking.

## Availability of supporting data

All raw and analyzed data is available on GigaDB at http://gigadb.org/site/index. All files and data are distributed under the Creative Commons CC0 waiver, with a request for attribution. The data are organized into 8 main folders for the 4 different data sets (see Table 4).

**Table 4.**
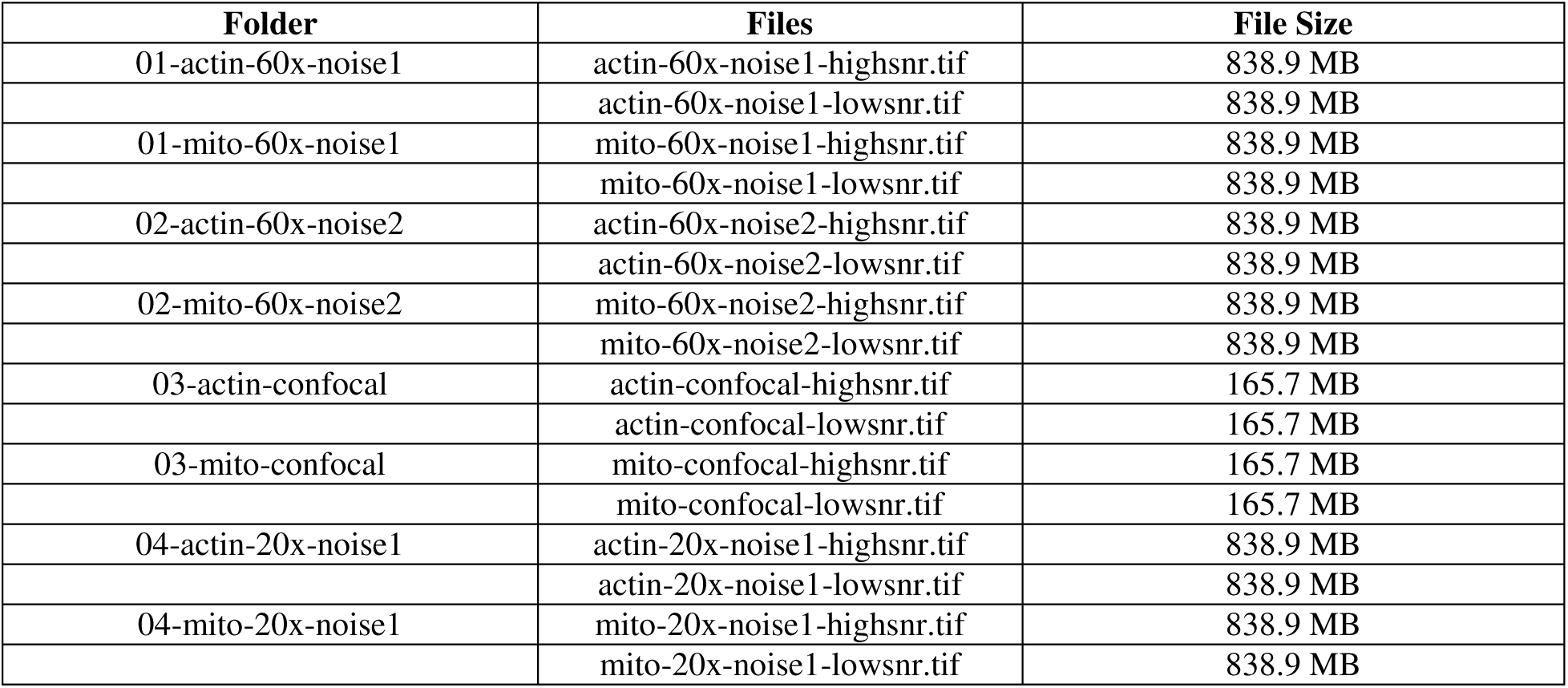
Description of the data files.

| Folder | Files | File Size |
| --- | --- | --- |
| 01-actin-60x-noise1 | actin-60x-noise1-highsnr.tif | 838.9 MB |
|  | actin-60x-noise1-lowsnr.tif | 838.9 MB |
| 01-mito-60x-noise1 | mito-60x-noise1-highsnr.tif | 838.9 MB |
|  | mito-60x-noise1-lowsnr.tif | 838.9 MB |
| 02-actin-60x-noise2 | actin-60x-noise2-highsnr.tif | 838.9 MB |
|  | actin-60x-noise2-lowsnr.tif | 838.9 MB |
| 02-mito-60x-noise2 | mito-60x-noise2-highsnr.tif | 838.9 MB |
|  | mito-60x-noise2-lowsnr.tif | 838.9 MB |
| 03-actin-confocal | actin-confocal-highsnr.tif | 165.7 MB |
|  | actin-confocal-lowsnr.tif | 165.7 MB |
| 03-mito-confocal | mito-confocal-highsnr.tif | 165.7 MB |
|  | mito-confocal-lowsnr.tif | 165.7 MB |
| 04-actin-20x-noise1 | actin-20x-noise1-highsnr.tif | 838.9 MB |
|  | actin-20x-noise1-lowsnr.tif | 838.9 MB |
| 04-mito-20x-noise1 | mito-20x-noise1-highsnr.tif | 838.9 MB |
|  | mito-20x-noise1-lowsnr.tif | 838.9 MB |

## Abbreviations

SSIM: structural similarity index
PSNR: peak signal to noise ratio
NA: numerical aperture

## Ethics approval and consent to participate

Not applicable

## Consent for publication

Not applicable

## Author Contributions

TN: analyzed data

JB: acquired data

RM: acquired data

TK: analyzed data

JV: conceived project, analyzed data, supervised research, wrote the paper

GH: conceived project, acquired data, analyzed data, supervised research, wrote the paper

